# Fatty acid auxotrophy as driver of Lactobacillus symbiosis in the female urinary tract

**DOI:** 10.64898/2026.04.23.720460

**Authors:** Michael L. Neugent, Hrishikesh H. Dalvi, Ceejay N. Saenz, Jessica L. Gauch, Nikki S. Koonjbearry, Jessica Komarovsky, Katherine C. Lim, Eve Spiro, Nicole J. De Nisco

## Abstract

The female urinary microbiome (FUM) has emerged a promising therapeutic target for recurrent urinary traction infection (rUTI). The FUM is dominated by a phylogenetically coherent group of *Lactobacillus* species whose metabolic relationship with the host remains poorly understood. Here, we demonstrate that fatty acid (FA) auxotrophy is a universal, conserved trait of FUM *Lactobacillus* species, mechanistically underpinned by complete loss or inactivation of the fab operon encoding Type II fatty acid synthesis (FASII). Stable isotope tracing with [U-¹³C] glucose confirms that *L. crispatus* and *L. gasseri* incorporate no glycolysis-derived carbon into cellular lipids, while generalist, non-FUM Lactobacillaceae species retain *de novo* FA biosynthesis. Gene synteny analysis across the Lactobacillaceae family reveals that fab operon loss is strongly associated with host-adapted lifestyles. Despite phenotypic FA auxotrophy among *L. crispatus* strains, a complete syntenic fab operon was identified in 31.9% of *L. crispatus* genomes. However, genomic inspection revealed a pervasive, inactivating frameshift mutation in fabH, the gene encoding the rate-limiting initiation condensing enzyme, in fab+ *L. crispatus* genomes highlighting two distinct evolutionary pathways of FA auxotrophy among this critically important FUM species. FA specificity assays establish *L. crispatus and L. gasseri* have an obligate requirement for monounsaturated FAs of C14–C20 chain length and that these species do not structurally modify supplemented FAs. Metabolic tracing further demonstrates that FUM lactobacilli directly scavenge and incorporate FAs from human bladder epithelial lipids, producing a bacterial membrane composition that mirrors the host. These findings establish FA auxotrophy as a defining adaptation to the female urogenital niche, with direct implications for development of microbiome-based therapies for rUTI.

## Introduction

The human microbiome comprises taxonomically and functionally distinct communities across anatomical sites whose composition is shaped by the biochemical constraints of their respective niches.^1^ A defining feature of host-adapted commensals is adaptive gene loss: repeated cycles of reductive evolution produce organisms whose biosynthetic repertoires are calibrated to the nutrient landscape of a particular host environment.^2–4^ This phenomenon, well-documented among obligate intracellular bacteria, increasingly emerges as a broader principle governing niche specialization among host-associated microbiota.^5^ Understanding the metabolic dependencies that underpin niche adaptation is therefore central to interpreting microbiome function, colonization requirements and to designing microbiome-based interventions.

The female urogenital microbiome (FUM) represents one of the most compositionally distinct microbial communities in the human body. Unlike the gut or oral cavity, where high alpha-diversity is associated with health, a *Lactobacillus*-dominated low-diversity state defines a healthy vaginal microbiome and, more recently, urinary microbiome in reproductive-age women.^1,6–8^ This unique composition, largely absent in other species, including non-human primates, reflects a co-evolutionary relationship between specific *Lactobacillus* lineages and the human host that is absent in other mammals.^9,10^ Population-scale surveys, including the ISALA cohort, reveal that FUM *Lactobacillus* species are distributed unevenly across ethnicities and geographic regions, implying that this host-microbe relationship is under continued co-evolutionary pressure.^11^ The dominant FUM *Lactobacillus* species, *L. crispatus*, *L. gasseri*, *L. jensenii*, *L. iners*, and *L. johnsonii*, form a phylogenetically coherent group whose lifestyle diverges markedly from generalist *Lactobacillaceae* such as *Lacticaseibacillus rhamnosus* or *Lactiplantibacillus plantarum*. High urogenital abundance of *L. crispatus* in particular is associated with protection against bacterial vaginosis (BV), urinary tract infection (UTI), and sexually transmitted infections including HIV acquisition, while dysbiotic states characterized by loss of *L. crispatus* dominance confer substantially elevated disease risk.^12–14^ Protective mechanisms of FUM lactobacilli are partially attributed to lactic acid production, which maintains acidic vaginal pH, and to the production of antimicrobial peptides and biosurfactants.^15–19^ Despite their importance to female urogenital health, the molecular underpinnings of FUM *Lactobacillus* fitness within the urogenital niche remain incompletely understood.

The clinical significance of FUM *Lactobacillus* species has motivated substantial interest in probiotic and live biotherapeutic product (LBP) development for urogenital indications.^20–24^ BV affects approximately 23-29% of reproductive-age women worldwide and is the most prevalent vaginal syndrome globally, yet recurrence rates following standard antibiotic therapy exceed 50% at 12 months.^25,26^ UTI, affecting 400 million people annually, similarly suffers from high recurrence and rising antimicrobial resistance especially in postmenopausal populations which tend to be depleted in urogenital lactobacilli .^8,27–30^ Microbiome-based therapies offer a compelling alternative to restore or reinforce a protective *Lactobacillus*-dominant urogenital state.^7^ A central obstacle; however, is achieving stable colonization of the target niche. Indeed, *L. crispatus* probiotic therapeutic failure for recurrent UTI, has been attributed to poor colonization.^31^ Administered probiotic strains must compete with established microbiota and persist within an environment whose physicochemical and nutritional parameters they must be able to exploit.^32,33^ Rational probiotic design therefore demands a quantitative understanding of the metabolic relationship between FUM lactobacilli and the host environment. Current formulation and delivery strategies for FUM modulation largely lack this mechanistic grounding in the chemical underpinnings of the human urogenital tract and microbiome interaction.

Relative to gut commensals, the nutrient requirements of FUM lactobacilli are poorly characterized. The pathobiont, *L. iners* has a well-documented requirement for multiple amino acids and nucleotide precursors, consistent with extensive host dependency.^34,35^ Metabolic dependencies are increasingly recognized as a defining feature of host-adapted bacteria, reflecting the selective advantage of offloading biosynthetic costs onto a metabolite-replete host environment.^2,36^ For other FUM species, metabolic characterization has been sparse. One study established that the fatty acids (FAs) oleic acid (OA) and 10-hydroxystearic acid promote *L. crispatus, L. gasseri, L. jensenii, and L. mulleris* growth, but inhibit the BV-associated *L. iners* growth. This growth stimulation was attributed to the presence of the genes *ohyA,* encoding an oleate hydratase, and *farE,* encoding a FA efflux pump, in *L. crispatus, L. gasseri, L. jensenii, and L. mulleris* that are absent in *L. iners.*^37^ These genes are required for conversion of OA to 10-hydroxystearic acid and for OA resistance, respectively. Importantly, the authors found that OA promotes *L. crispatus* dominance in an in vitro BV model.^37^ This study also noted that OA stimulated the growth of FUM species *L. crispatus, L. gasseri, L. jensenii, and L. mulleris* in lipid-depleted, but not FA-free, undefined MRS+CQ medium suggesting a potential FA auxotrophy. The FA biosynthesis precursor, acetate, did not stimulate growth; however, these experiments did not conclusively demonstrate auxotrophy as they lacked a species capable of de novo FA biosynthesis as an essential positive control.^37^ Furthermore, the prevalence of this potential FA requirement across the *Lactobacillus* genus and broader Lactobacillaceae family was not explored. Therefore, whether FA auxotrophy represents a genuine and conserved metabolic hallmark of FUM lactobacilli, and if so, its molecular basis, has remained unresolved.

Both bacteria and plants primarily utilize the Type II fatty acid synthesis (FASII) pathway, encoded by the *fab* operon, to generate essential fatty acid building blocks for membrane phospholipids.^38^ The pathway initiates with the acetyl-CoA carboxylase-mediated carboxylation of acetyl-CoA to malonyl-CoA, which is subsequently transferred to acyl carrier protein (ACP) by FabD. Chain initiation is catalyzed by the β-ketoacyl-ACP synthase FabH, which condenses acetyl-CoA with malonyl-ACP to generate the first 4-carbon intermediate. Iterative two-carbon elongation cycles, involving the reductases FabG and FabI and the dehydratase FabZ, extend the acyl chain to the physiological lengths required for membrane phospholipid synthesis.^39,40^ Unsaturated fatty acids are generated in Gram-positive bacteria through an anaerobic pathway in which FabA or the FabZ-like dehydratase introduces a *cis*-double bond during elongation, yielding monounsaturated products including palmitoleic acid (16:1Δ11*cis*) and vaccenic acid (18:1Δ11*cis*).^41^ As lipids are one of the four essential building blocks of life, instances of complete FA auxotrophy are relatively rare across the bacterial kingdom.^42^

Here, we investigate the hypothesis that FUM *Lactobacillus* species are obligate FA auxotrophs that have undergone lineage-specific loss of the FASII pathway. To test this hypothesis, we pursued three interconnected goals: (1) to phenotypically characterize FA requirements across a diverse panel of *Lactobacillaceae* species using defined growth media and stable isotope tracing; (2) to determine the phylogenetic distribution and genomic basis of FA auxotrophy through comparative analysis of *fab* operon synteny across the *Lactobacillaceae* family; and (3) to define the repertoire of FAs utilized by FUM *Lactobacillus* species and test the hypothesis that these species can directly utilize host lipids to fulfill their exogenous FA requirement. This work expands the foundational metabolic characterization of FUM lactobacilli and extends phylogenomic frameworks characterizing lifestyle diversity within the *Lactobacillaceae*.

We report that FA auxotrophy is a phylogenetically conserved trait of FUM lactobacilli, mechanistically underpinned by widespread loss or inactivation of the FASII *fab* operon. Using [U-^13^C]glucose stable isotope tracing, we demonstrate that *L. crispatus* and *L. gasseri* incorporate zero glycolysis-derived carbon into cellular lipids, while generalist *Lacticaseibacillus* and *Lactiplantibacillus* species maintain active *de novo* FA biosynthesis. Comparative genomics of 123 *Lactobacillaceae* genomes reveals that *fab* operon loss is strongly associated with host adaptation, particularly within the genus *Lactobacillus* but also with the genera *Limosilactobacillus, Apilactobacillus,* and *Companilactobacillus*. Contrary to a prior report, we found that 31.91% of *L. crispatus* strains have retained full syntenic *fab* operons, despite having a phenotypic exogenous FA dependency. Within 93.81% of these Fab+ *L. crispatus* strains, we identify a highly conserved inactivating mutation in the FASII initiation enzyme, FabH providing an evolutionary snapshot into *L. crispatus* metabolic host adaptation. FA specificity assays establish that *L. crispatus* and *L. gasseri* FA requirement is fulfilled by monounsaturated FAs, with optimal growth requiring C16–C18 chain lengths and that saturated FAs cannot serve as sole lipid sources. Finally, using metabolic tracing experiments with labeled bladder epithelial cell lipids, we find that *L. crispatus* and *L. gasseri* directly incorporate fatty acids from host lipids into their membranes resulting in a bacterial FA membrane composition nearly identical of that of the host. These findings define FA auxotrophy as an evolutionary adaptation of FUM lactobacilli to the lipid-replete urogenital niche and establish host-derived FAs as a fundamental metabolic axis governing colonization fitness and bacterial membrane FA composition, with direct implications for probiotic development and microbiome-aware strategies for the management of recurrent UTI in women.

## Results

### Female urogenital microbiome (FUM) lactobacilli require an exogenous fatty acid source

To begin to understand the nutrient requirements of urogenital lactobacilli, we first sought to investigate the impact of Tween-80, a component of the De Man, Rogosa, and Sharpe (MRS) medium that is typically used to cultivate Lactobacillaceae, in Lactobacillaceae species growth phenotypes. We screened a diverse panel of species from the Lactobacillaceae family in a modified De Man, Rogosa, and Sharpe (mMRS) medium containing all standard MRS ingredients with the exception of Tween-80 as an experimental condition. In this screen, many Lactobacillaceae species exhibited a strict growth reliance on Tween-80. This requirement for Tween-80 was universal among all female urogenital microbiome (FUM)-associated species, including *L. crispatus*, *L. gasseri*, *Lactobacillus paragasseri*, *L. johnsonii*, *L. jensenii*, *Limosilactobacillus colehominis*, *Limosilactobacillus vaginalis*, and *Limosilactobacillus portuensis* (**Fig. 1A**). In contrast, more generalist species such as *L. rhamnosus*, *L. plantarum*, *Lacticaseibacillus casei*, *Lacticaseibacillus paracasei*, *Limosilactobacillus fermentum*, and *Lacticaseibacillus zeae* achieved robust growth in the absence of Tween-80 (**Fig. 1A**). Interestingly, *Lactobacillus acidophilus*, a common inhabitant of the oral cavity and gastrointestinal tract also required Tween-80 for growth. The species *Lactobacillus delbrueckii*, *Levilactobacillus brevis*, and *Ligilactobacillus salivarius* did not require Tween-80 for growth in mMRS but did demonstrate further growth stimulation upon Tween-80 addition.

**Figure 1.**
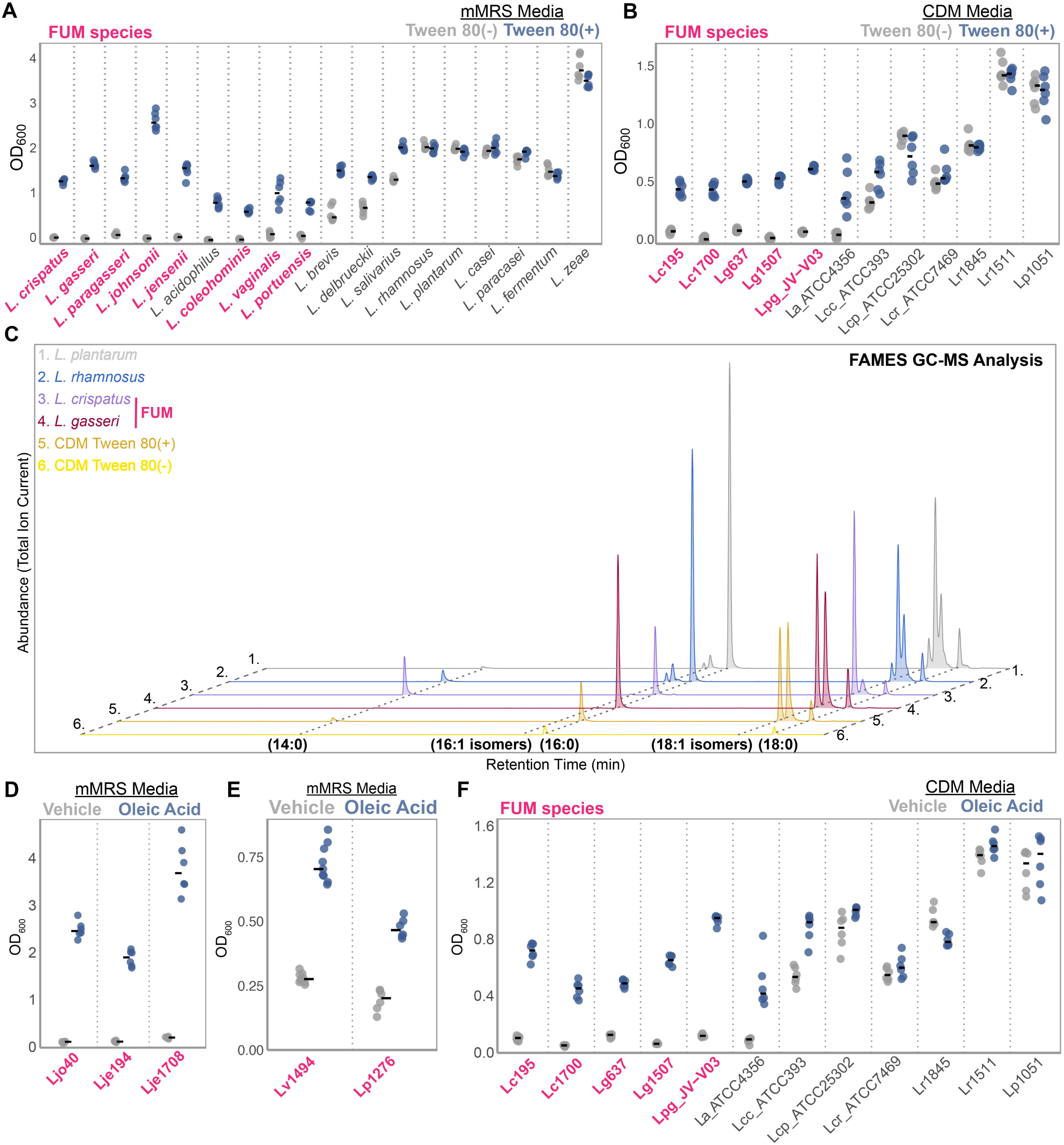
Female urogenital microbiome (FUM) lactobacilli require an exogenous fatty acid source. (A,B) End-point growth assays (OD_600_) of various members of *Lactobacillaceae* family cultured in **(A)** modified DeMan, Rogosa, Sharpe lactobacilli growth medium (mMRS) and **(B)** chemically defined medium (CDM) with and without Tween80. Female urogential microbiome (FUM) species are colored pink. **(C)** Gas chromatography-mass spectrometry (GC-MS) chromatograms of fatty acid methyl esters (FAMEs) of *Lactobacillaceae* family members after grown in presence of Tween80. **(D,E)** Endpoint growth assay (OD_600_) of FUM isolates of *L. johnsonii*, *L. jensenii*, *L. vaginalis* and *L. portuensis* species in modified MRS (mMRS) supplemented with oleic acid or BSA vehicle. **(F)** Endpoint growth assay (OD_600_) of FUM and non-FUM isolates, in chemically defined medium (CDM) supplemented with BSA (vehicle) or oleic acid conjugated with BSA.

We further validated this phenotype using a completely chemically defined media (CDM) originally designed for experimentation with streptococci.^43,44^ Consistent with the mMRS results, FUM species, including *L. crispatus* (Lc195, Lc1700), *L. gasseri* (Lg637, Lg1507), and *L. paragasseri* (Lp_JV-V03), demonstrated strict dependence on Tween-80 for growth in CDM while generalist *Lactiplantibacillus* and *Lacticaseibacillus* species grew in CDM independently of Tween-80 (**Fig. 1B**). FUM species *L. johnsonii*, *L. jensenii*, *L. colehominis*, *L. vaginalis*, and *L. portuensis* could not be tested in CDM because they do not grow in CDM even with Tween-80 supplementation suggesting additional nutrient dependencies of these species.

Chemically, Tween-80 is predominantly composed of oleic acid (OA; 18:1.49*cis*), among other FAs, conjugated to a polar polysorbate headgroup. We hypothesized that Tween-80 serves as a source of fatty acids (FAs) and that the FUM lactobacilli may require Tween-80 because they are FA auxotrophs. Gas chromatography-mass spectrometry (GC-MS) analysis of fatty acid methyl esters (FAMEs) confirmed Tween-80 contained abundant oleic (18:1.49*cis*) acid as well as myristic (14:0), palmitic (16:0), elaidic (18:1.49*trans*) and stearic acid (18:0) (**Fig. 1C**). FAMES analysis of extracted bacterial lipids revealed that the cellular fatty acid profiles of FUM lactobacilli, *L. crispatus*, *L. gasseri,* qualitatively mirrored the FA composition of the CDM + Tween-80 media. Conversely, the FAMEs profiles of generalist species, *L. rhamnosus* and *L. plantarum* included FAs not detected in the media, such as 16:1 isomers and an additional 18:1 isomer (**Fig. 1C**). These data support the hypothesis that, unlike generalist *Lactobacillaceae* species, FUM lactobacilli may require Tween-80 to fulfill a specific FA nutrient requirement.

To directly test if the Tween-80 dependency was specific to FAs, we supplemented oleic acid, the most abundant FA in Tween-80, into liquid cultures lacking Tween-80 using bovine serum albumin (BSA) as a solubilization vehicle.^45^ For FUM species requiring mMRS as a basal medium, such as *L. johnsonii*, *L. jensenii*, *L. vaginalis*, and *L. portuensis*, we observed robust growth only in OA-supplemented mMRS relative to the empty BSA vehicle control (**Fig. 1D, E**). The same OA-supplementation experiments were performed for *L. crispatus*, *L. gasseri*, and *L. paragasseri* in the cleaner, defined CDM and we observed a strict requirement for an exogenous FA source, whereas generalist species such as *L. casei*, *L. paracasei*, *L. rhamnosus*, and *L. plantarum* grew robustly in the absence of supplementation (**Fig. 1F**). Collectively, these data suggest that urogenital lactobacilli are unable to sustain growth without environmental **FAs**.

### FUM lactobacilli cannot synthesize fatty acids from glucose

The Type II fatty acid synthesis (FASII) pathway, initiated by the conversion of acetyl-CoA to malonyl-CoA, is the canonical route for membrane lipid biogenesis in bacteria.^39^ Based on the strict exogenous FA requirement observed among urogenital-associated species in Figure 1, we hypothesized that FUM lactobacilli lack the metabolic capacity for *de novo* FA biosynthesis. To test this, we performed stable isotope tracing using [U-^13^C]glucose in CDM supplemented with Tween-80. We traced the flux of glycolysis-derived carbon into the cellular lipid pool using GC-MS analysis of FAMEs (**Fig. 2A**). In key FUM species, *L. crispatus* and *L. gasseri*, we observed no ^13^C incorporation into any identified FA species, including myristic (14:0), palmitic (16:0), stearic (18:0), palmitoleic (16:1), oleic (18:1.49*cis*), or elaidic (18:1.49*trans*) acids (**Fig. 2B, C**). The mass spectra displayed only the unlabeled (M+) isotopologues, confirming that these cellular lipids were derived entirely from the exogenous Tween-80 source rather than central carbon metabolism.

**Figure 2.**
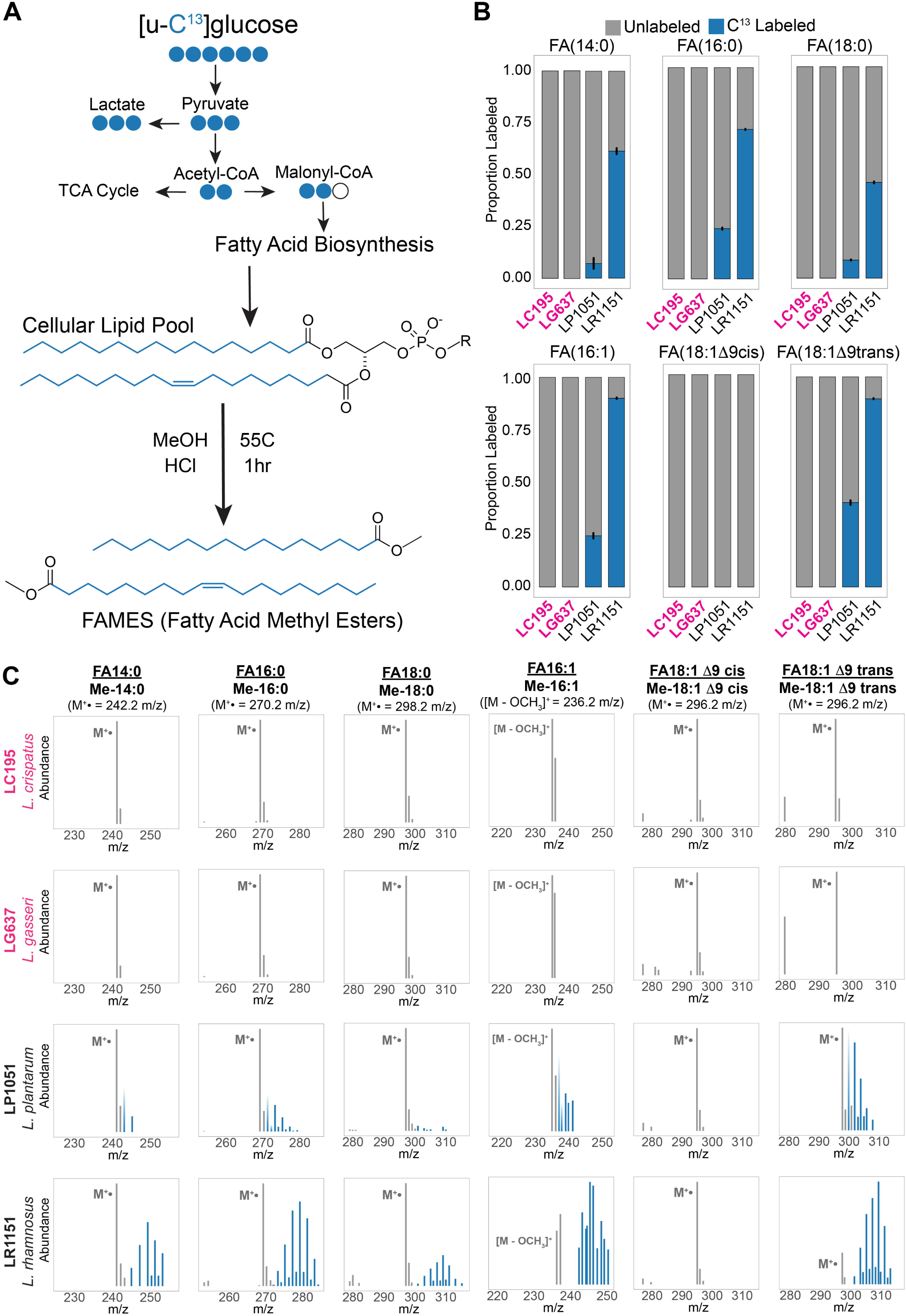
FUM lactobacilli cannot synthesize fatty acids from glucose. **(A)** Schematic overview of tracing stable isotope [U-^13^C] glucose in central carbon metabolism which following glycolysis to form pyruvate, which is converted to acetyl-CoA and finally enters fatty acid biosynthesis as malonyl-CoA. These newly formed FAs are incorporated into the cellular lipid pool and are derivatized into FAMEs for further GC-MS analysis. **(B)** Proportional enrichment of ¹³C-labeled (blue) versus unlabeled (grey) fatty acids across FUM strains (*L. crispatus* Lc195 and *L. gasseri* Lg637) and non-FUM strains (*L. rhamnosus* Lr1511 and *L. plantarum* Lp1051). **(C)** Mass spectra of targeted FAMEs ^13^C enrichment.

Conversely, the generalist species *L. plantarum* and *L. rhamnosus* exhibited robust ^13^C enrichment across their cellular FA profiles, confirming active *de novo* biosynthesis from glucose (Fig. 2B, C). Notably, while generalist Lactobacillaceae species synthesized labeled saturated FAs from glucose, they did not incorporate ^13^C into oleic acid (18:1.49*cis*). Instead, the labeled carbon was directed into the synthesis of the 18:1.49*trans* isomer, elaidic acid and membrane oleic acid was unlabeled (**Fig. 2B**). This distinction reveals that while the generalist species *L. plantarum and L. rhamnosus* possess the machinery to synthesize specific monounsaturated FAs, they also scavenge the abundant 18:1.49*cis* provided by Tween-80 while biosynthesizing the *trans*-isomer. Collectively, these data provide strong biochemical evidence that FUM lactobacilli lack the metabolic capacity to convert glycolysis-derived carbon into fatty acids.

### Lack of the Fab operon underpins FUM lactobacilli fatty acid auxotrophy

To determine the genetic basis of the observed FA auxotrophy among FUM lactobacilli, we performed a comparative genomic analysis of the Fab operon, which encodes Type II fatty acid synthesis (FASII) pathway, across the Lactobacillaceae family using publicly available, high-quality genomes with low fragmentation.^39^ With strict quality-control criteria, we assembled a dataset of 123 representative family-level genomes sampled across lifestyles ranging from environmental Free-living (n= 40), Host-adapted (n= 42), and Nomadic (n= 13) representing 31 genera.^46^ We reconstructed the phylogeny of representative Lactobacillaceae species based on average nucleotide identity (ANI), and used gene synteny analysis to annotate the presence of a complete Fab operon containing the required gene set for long chain FA biosynthesis (**Fig. 3A, B, C**).^39^ Our synteny analysis revealed a lineage-specific, widespread loss of the FASII pathway that correlates strongly with host adaptation (*P_Fisher-Exact_ =* 5.1x10^-5^) (**Fig. 3A, B**). Among the analyzed Lactobacillaceae species, the Fab operon was universally present in 70 species (56.91%), completely absent in 40 species (32.52%), and heterogeneously present in 13 species (10.56%). While nomadic and free-living generalist taxa, such as *Lacticaseibacillus* (*L. casei*, *L. rhamnosus*) and *Lactiplantibacillus* (*L. plantarum*), universally retain the Fab operon, the genus *Lactobacillus* exhibits a widespread absence of these biosynthetic genes (**Fig. 3A, B**).^46^ This genus encompasses the dominant FUM species, including *L. crispatus*, *L. gasseri*, *L. jensenii*, *L. johnsonii* and *L. iners*, suggesting that the loss of FA biosynthesis may be a phylogenetically conserved trait among urogenital lactobacilli. Interestingly, three FUM species, *L. crispatus*, *L. vaginalis,* and *L. portuensis* exhibited heterogeneous Fab operon conservation genotypes with 31.10% (93/299), 14.28% (5/35), and 31.25% (5/16) of publicly available genomes possessing the Fab operon, respectively (**Fig. 3A, B**).

**Figure 3.**
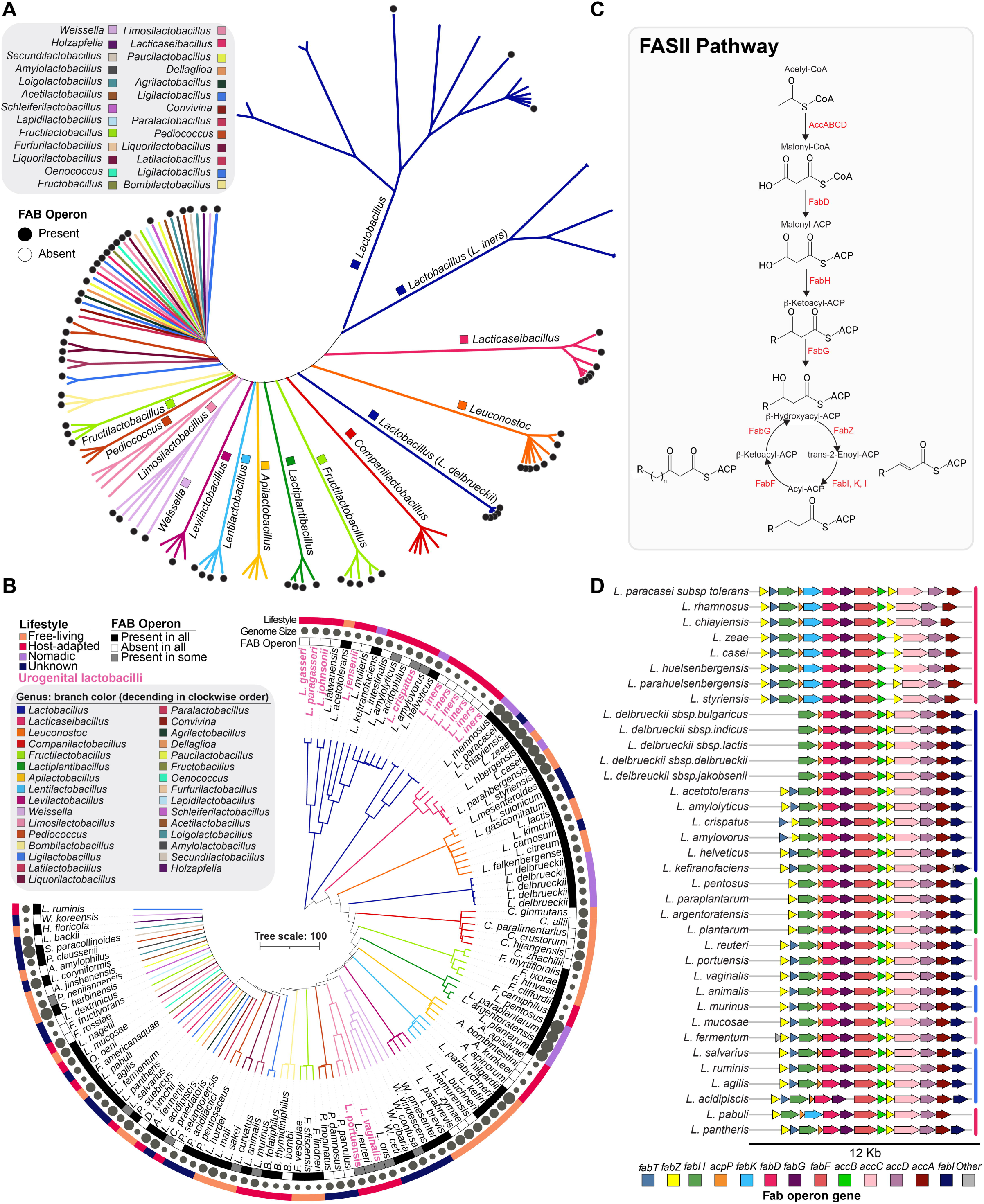
Fab operon conservation and synteny across Lactobacillaceae. (A) Unrooted and (B) rooted family-level phylogenetic dendrograms of 123 representative Lactobacillaceae genomes based on average nucleotide identity (ANI), with branches colored by genus. In (B), urogenital lactobacilli are labeled in pink; Fab operon status is indicated by black (present in all strains), grey (present in some strains), or white (absent in all strains) squares; genome size is indicated by grey circles; and lifestyle categories are color-coded as free-living (orange), host-adapted (pink), nomadic (purple), or unknown (blue). (C) Schematic of the FASII pathway with the enzymatic role of each Fab operon gene indicated. (D) Comparative synteny map of Fab operons extracted from 36 Lactobacillaceae species, with each coding sequence depicted as a directional arrow colored by gene identity and operons aligned to fabF.

In both the unrooted and rooted trees, there was considerable genetic divergence of two *Lactobacillus* species, *L. iners* and *L. delbrueckii,* from the rest of the species in the genus as they formed separate branches within the family level tree (**Fig. 3A, B**). The Fab operon was universally absent from FUM *L. iners* but universally present in non-FUM *L. delbreuckii.* Detailed synteny analysis of Fab operons among the Lactobacillaceae family showed distinct variable operon structures between species (**Fig. 3D)**. The majority of *Lacticaseibacillus* species encode a complete, syntenic Fab operon containing two copies of *fabZ* (dehydratase), *fabT* (transcriptional regulator), *fabH* (initiation condensing enzyme), *acp* (acyl carrier protein), *fabK* (enoyl-ACP reductase), *fabD* (malonyl-CoA-ACP transacylase), *fabG* (reductase), fabF (elongation condensing enzyme), *fabZ* (dehydratase), and the *accABCD* genes (acetyl-CoA carboxylase complex subunits) with the exception of *Lacticaseibacillus pabuli* which lacks *fabT* and *Lacticaseibacillus pantheris* which encodes the FabI type enoyl ACP-reductase instead of FabK (**Fig. 3D**). *Lactobacillus, Limosilactobacillus,* and *Ligilactobacllus spp.* have a similar operon structure, but universally encode FabI instead of FabK, and *Lactobacillus delbrueckii* species notably lack a second copy of *fabZ* and *fabT. Lactiplantibacillus* species also encode FabI instead of FabK and lack the regulatory gene *fabT* (**Fig. 3D**).

### A widespread nonsense mutation inactivates the FASII pathway in Fab(+) *L. crispatus*

The observation that 31.10% of *L. crispatus* genomes possess the Fab operon while our data suggest the species is phenotypically auxotrophic prompted us to investigate the genetic architecture of the Fab+ *L. crispatus* strains. We reconstructed the *L. crispatus* phylogeny using 304 high-quality publicly available genomes downloaded from NCBI RefSeq by excluding atypical genomes and metagenome-assembled genomes and selecting those with < 200 contigs and >74.12% CheckM completeness. The genomes submitted with geographic location data were from 15 different countries. The majority of the *L. crispatus* genomes obtained were from the USA (n= 177, 57.89%), followed by Italy (n= 49, 16.11%), South Africa (n= 27, 8.88%), India (n= 10, 3.28%), China (n= 7, 2.30%), Brazil (n= 4, 1.31%), the United Kingdom (n= 3, 0.98%), South Korea (n= 2, 0.65%), France (n= 2, 0.65%), Germany (n= 2, 0.65%), Japan (n= 2, 0.65%), Ireland (n= 2, 0.65%), Russia (n= 1, 0.32%), Finland (n= 1, 0.32%), and Turkey (n= 1, 0.32%). Additionally, 14 genomes were submitted with no geographical location data (Unconfirmed, n=14, 4.60%). Our analysis identified three genetically distinct clades with Clade 1 being strongly enriched in strains isolated from the GI tract with (58/99) 58.58% GI, followed by (41/99) 41.41% vaginal, (12/99) 12.12% urine, (4/99) 4.04% oral, ocular (1/99) 1.01% and (10/99) 10.10% from unconfirmed sources. Clade 2 was mostly composed of vaginal isolates (70/77) 90.90%, followed by urine (5/77) 6.49%, GI tract (1/77) 1.29% and unknown (1/77) 1.29%. Clade 3 was mainly comprised of vaginal (123/128) 96.09%, and urine isolates (5/128) 3.90% (**Fig. 4A**). Clades 1 and 2 exhibited very limited Fab operon presence with (6.06% (6/99) and 1.29% (1/77) of strains encoding the operon, respectively). In contrast, Clade 3 was strongly enriched for the Fab operon, with 70.31% (90/128) of strains being genotypically Fab+ (**Fig. 4B**).

**Figure 4.**
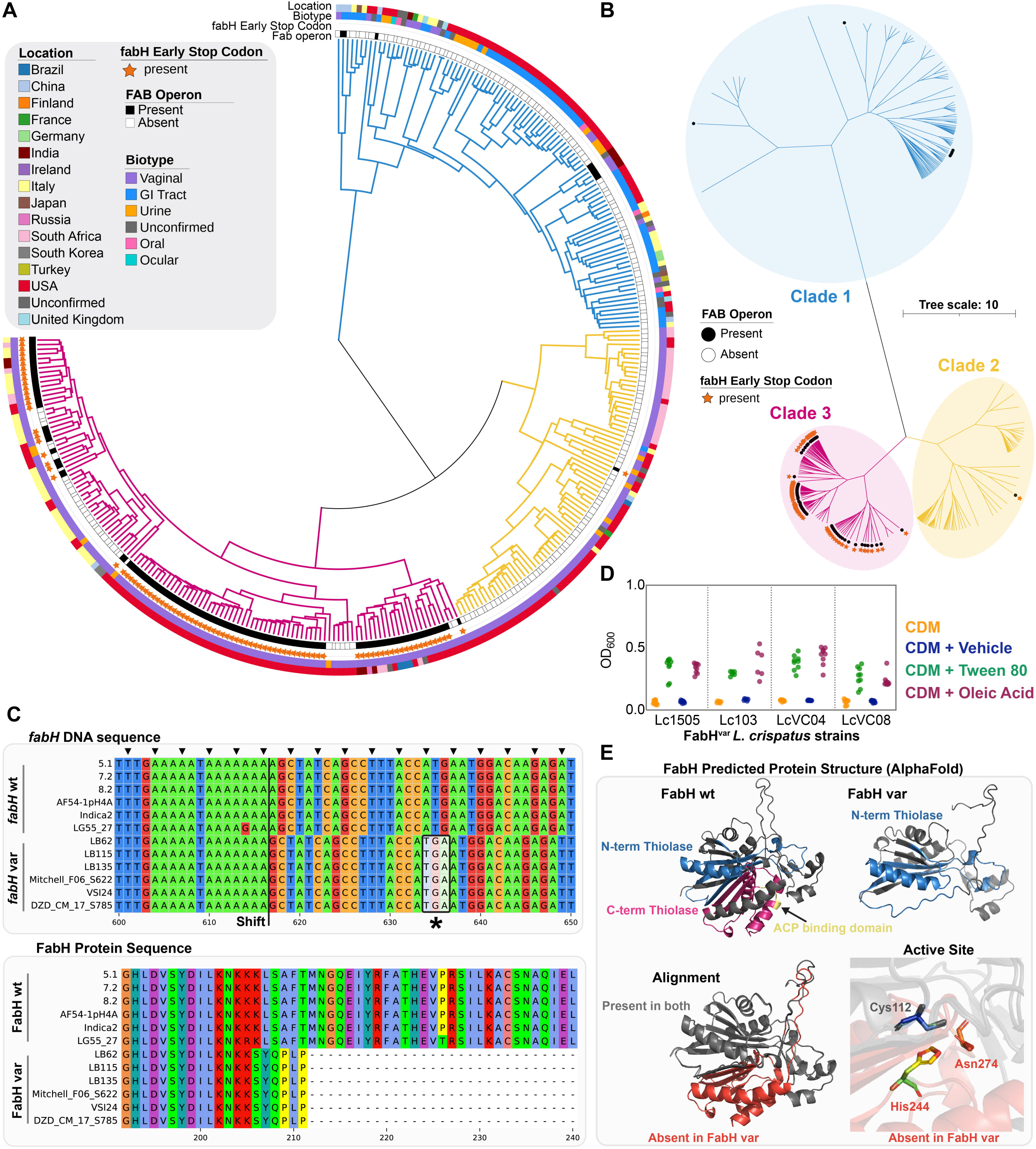
Fab operon variation and fabH inactivation in *Lactobacillus crispatus*. **(A)** Rooted and **(B)** unrooted ANI-based phylogenetic dendrograms of 304 high-quality L. crispatus genomes, with three main clades distinguished by color (blue, yellow, pink). Fab operon presence is indicated by a black square/circle and a premature stop codon in fabH by a gold star. In (A), strain isolation source (biotype) and geographic origin are indicated by categorical color. **(C)** Nucleotide (top) and amino acid (bottom) sequence alignments of wild-type fabH and fabH variant sequences extracted from *L. crispatus* strains, with the premature stop codon position marked by a black asterisk. **(D, E)** AlphaFold3-predicted three-dimensional structures of wild-type FabH and the truncated FabH variant.

We next examined the coding sequences of individual Fab operon genes in the Fab+ *L. crispatus* strains. We identified a pervasive mutation in *fabH*, the gene encoding the b-ketoacyl-ACP synthase III, which catalyzes the rate-limiting condensation of acetyl-CoA with malonyl-ACP to initiate fatty acid synthesis.^39,47^ This variant was present in 93.81% (91/97) of Fab+ *L. crispatus* strains, and more specifically in 100% of Fab+ clade 2 and 3 strains but in none of the clade 1 strains (**Fig 4A,B**). Alignment of wild-type and variant *fabH* sequences revealed a single nucleotide deletion within a poly-adenosine tract (nucleotides 610-617). This deletion causes a frameshift that introduces a premature stop codon (TGA) at nucleotides 634-636 (**Fig. 4C**). The resulting truncated FabH variant is predicted to be 210 amino acids long, compared to the 322 amino acids of the wild-type protein (**Fig. 4C**). To validate the functional impact of this mutation, we confirmed that FabH-variant strains (Lc1505, LC103, Lc_VC04, Lc_VC08) are phenotypic FA auxotrophs, characterized by a lack of growth in CDM without Tween-80 that is rescued by OA supplementation (**Fig 4D**). Structural prediction using AlphaFold revealed that the mutant protein lacks the C-terminal catalytic core essential for enzymatic activity, including the critical active site residues, His244 and Asn274 (**Fig. 4E**).^48–50^ These data suggest that even in *L. crispatus* lineages that have retained the Fab operon, a widespread inactivating mutation in the gene encoding the initiation enzyme, *fabH,* renders the pathway non-functional. These findings cement the idea that *L. crispatus* has evolved to rely on exogenous fatty acids with 98.02% of *L. crispatus* genomes being either Fab(-) or containing the inactive FabH variant genotype.

### *L. crispatus* and *L. gasseri* require exogenous unsaturated fatty acids

To further validate the idea that FUM lactobacilli rely on exogenous FAs, we characterized OA acid uptake for both FUM and non-FUM Lactobacillaceae using stable isotope tracing. Bacterial strains were cultured in CDM supplemented with [U-^13^C]oleic acid (18:1.49*cis*), and cellular FA pools were analyzed via FAMEs GC-MS. We observed a near-total mass shift to the ^13^C-labeled isotopologue (M^+^ 314 m/z) across all tested strains, including both FUM (*L. crispatus*, *L. gasseri*) and generalist (*L. rhamnosus*, *L. plantarum*) species, with negligible unlabeled OA (M^+^ 296 m/z) detected in the cellular FA pool (**Fig. 5A**). This indicates that generalist *L. rhamnosus* and *L. plantarum*, despite their de novo FA biosynthetic capabilities, do still uptake exogenous OA upon supplementation. Furthermore, as mass shifts were only observed for OA, our data suggest that neither FUM nor non-FUM Lactobacillaceae modify supplemented OA in a way that can be detected by the FAMES method. However, for FUM species this FA scavenging is not facultative but represents an obligate metabolic reliance, as they lack the capacity to synthesize alternative FAs.

**Figure 5.**
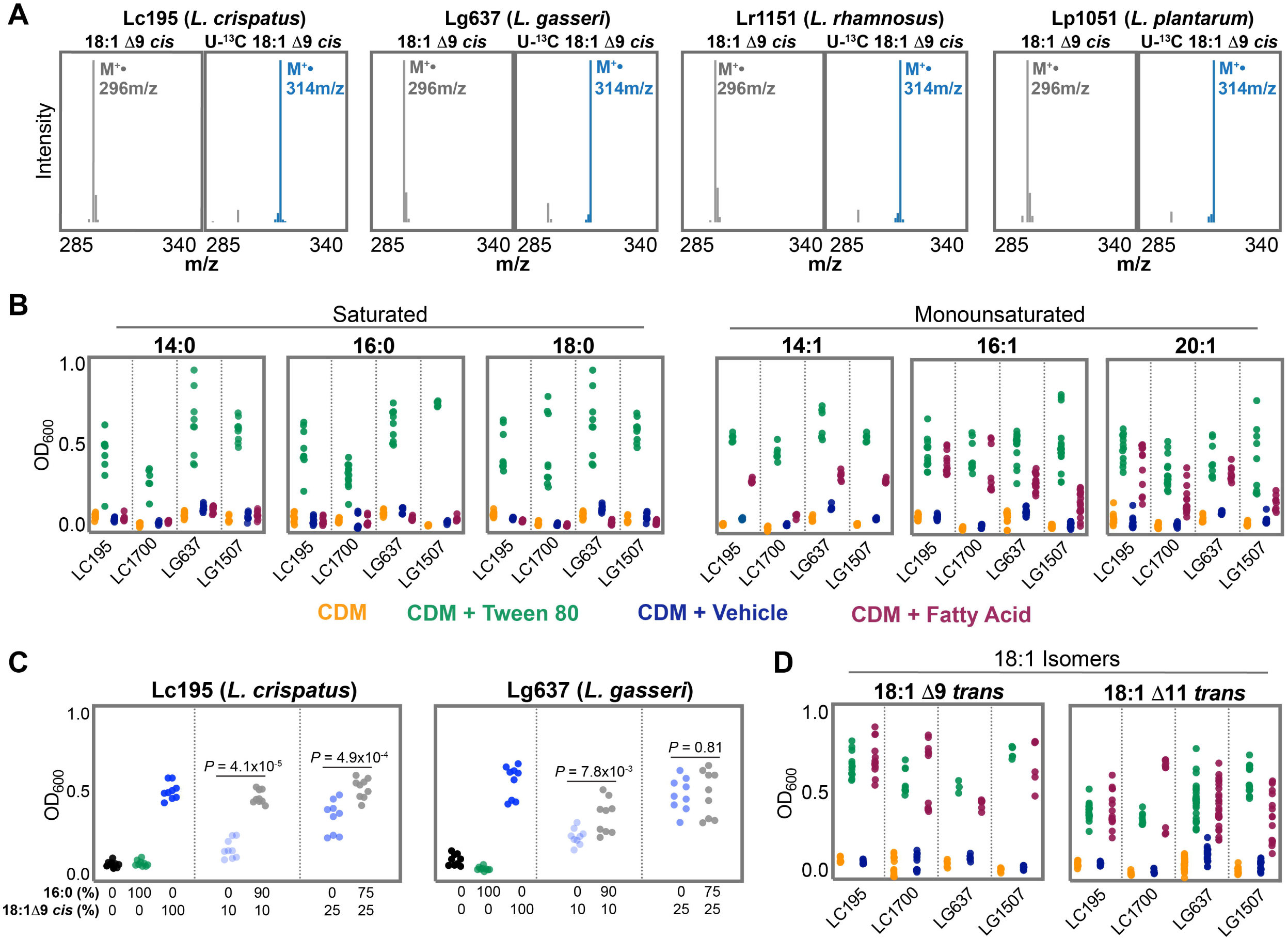
*L. crispatus* and *L. gasseri* require exogenous unsaturated fatty acids. **(A)** Mass spectra showing direct incorporation of exogenously supplemented unlabeled vs [U-^13^C]-labeled oleic acid into both FUM and non-FUM species. **(B)** Endpoint growth assays of two strains of *L. crispatus* (Lc195 and Lc1700) and *L. gasseri* (Lg637 and Lg1507) belonging to FUM-*Lactobacillus* in various saturated (14:0, 16:0, 18:0) and monounsaturated (14:1, 16:1 and 20:1) FAs. **(C)** Endpoint growth assay of FUM-*Lactobacillus* species in co-supplemented saturated (16:0, palmitic acid) and unsaturated (18:1Δ11cis, oleic acid) FAs in varying ratios. P-value generated via Wilcoxson Rank Sum test. **(D)** Endpoint growth assays of two strains of *L. crispatus* (Lc195 and Lc1700) and *L. gasseri* (Lg637 and Lg1507) in positional isomers of oleic acid (18:1Δ9 trans, elaidic acid and 18:1Δ11 trans, vaccenic acid).

To identify FA preferences, in terms of saturation and chain length, of prominent FUM species *L. crispatus* and *L. gasseri*, we supplemented CDM with individual saturated or unsaturated FA species. We observed that saturated FA, including myristic (14:0) and palmitic (16:0) acid, failed to support *L. crispatus* and *L. gasseri* growth as a sole FA source (**Fig. 5B**). In contrast, *L. crispatus* and *L. gasseri* were able to use monounsaturated FAs as a sole FA source, though with notable chain-length specificity. Myristoleic acid (14:1) supported moderate growth for both *L. gasseri* strains but only for one *L. crispatus* strain, Lc195. Palmitoleic acid (16:1) supported growth for all strains but not to the same levels as observed for Tween-80. Eicosenoic acid (20:1) also supported growth for most strains tested, but to a lesser extent for *L. gasseri* strain Lg1507 (**Fig. 5B**). These data suggest that saturated FAs cannot be utilized as a sole FA source *by L.* crispatus *and L. gasseri*, while FAs of chain length C14-C20 can solely support growth, but to different extents in different strains.

Although *L. crispatus* and *L. gasseri* cannot use saturated FAs as a sole FA source, our GC-MS FAMES experiments on these species grown in Tween-80, which also contains palmitic acid (16:0), suggested that the species do incorporate saturated palmitic acid (PA) when provided in a mixture (**Fig. 1C**). To conclusively determine if *L. crispatus* and *L. gasseri* can utilize saturated FAs in the presence of unsaturated FAs, we performed growth assays in CDM titrating with varying ratios of PA (16:0) to OA (18:1). *L. crispatus* Lc195 grew to a significantly higher density in 90/10 PA/OA acid than the 10% OA equivalent suggesting that PA is utilized by *L. crispatus* (**Fig. 5C**)*. L. gasseri* Lg637 exhibited more variable but statistically significant growth stimulation by PA in the 90/10 PA/OA condition compared to the 10% OA equivalent (**Fig. 5C**). These data demonstrate that although FUM lactobacilli possess an absolute requirement for exogenous unsaturated FAs, saturated FAs can also stimulate *L. crispatus* and *L. gasseri* growth when unsaturated FAs are also present.

Finally, we tested the stereochemical and positional specificity of the unsaturated FA requirement of predominant FUM species *L. crispatus* and *L. gasseri* as unsaturated FAs can also be found in the trans configuration and the double bond can occur at different positions in the carbon chain. Supplementation with naturally occurring C18 trans isomers, elaidic acid (18:1.49*trans*) and vaccenic acid (18:1.411*trans*), supported robust growth in *L. crispatus* strains Lc195 and Lc1700 as well as *L. gasseri* strains Lg637 and Lg1507 (**Fig. 5D**). These data indicate that the *L. crispatus* and *L. gasseri* exogenous FA metabolic requirement is satisfied by the presence of a double bond, independent of specific cis/trans geometry or .49/.411 positioning, and least for C18 FAs.

### Auxotrophic FUM lactobacilli can directly utilize host lipids as a FA source

Given the close mutualistic relationship between FUM lactobacilli and the human female urogenital tract, we hypothesized that FUM lactobacilli satisfy their strict FA requirement by scavenging lipids from their environment. In addition to their prominence in the healthy female vaginal microbiome *L. crispatus* and *L. gasseri* are two of the most predominant species of the healthy female urobiome.^28^ In the nutrient poor bladder environment, exfoliated bladder epithelial cells may serve as a potential FA source for bladder-resident FUM lactobacilli. To test if predominant FUM lactobacilli utilize bladder epithelial lipids as a FA source, we extracted total cellular lipids from two human bladder epithelial cell lines, 5637 and T24. We then assessed the capacity of these host-derived lipids to support bacterial growth as a sole FA source. Supplementation of the FA-free culture medium with lipid extracts from either 5637 or T24 cells was sufficient to restore robust growth in both *L. crispatus* LC195 and *L. gasseri* LG637, comparable to levels observed with Tween-80 (**Fig. 6A, B**).

**Figure 6.**
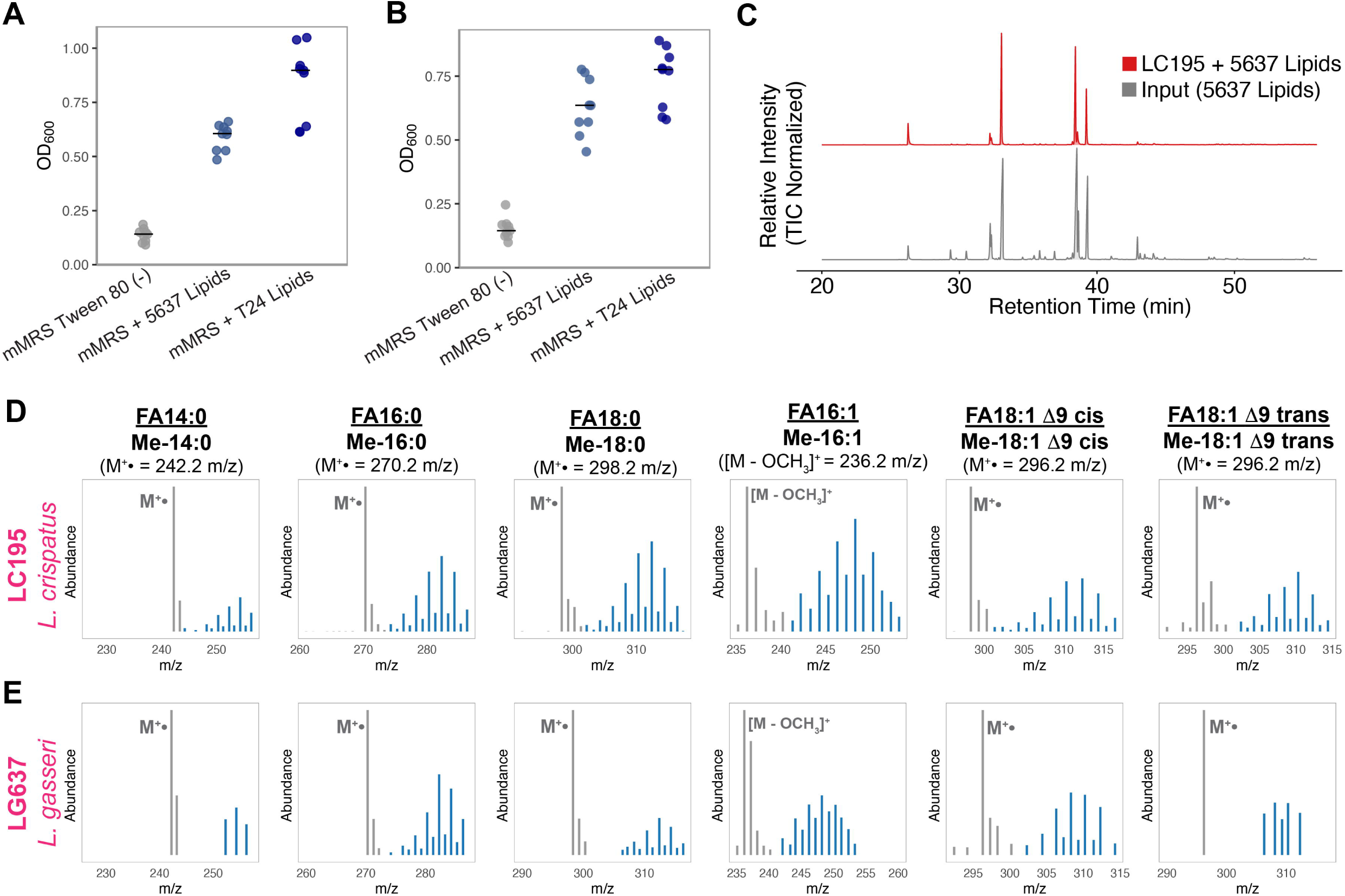
Auxotrophic FUM lactobacilli can directly utilize host lipids as a FA source. (A,B) End-point growth assay of representative Lc195(A) and Lg637(B) cultured in lipid-depleted modified De Man, Rogosa and Sharpe medium (mMRS Tween 80(-)). **(C)** GC-MS chromatograms depicting comparison of the total lipid profile of Lc195 supplemented with lipids extracted from 5637 cells vs only extracted 5637 lipids. **(E,F)** Mass spectra of various FAMEs from (E) *L. crispatus* Lc195 and (F) *L. gasseri* Lg637 grown in presence of ^13^C-labeled fatty acids from eukaryotic cells.

To evaluate how these host lipids influence the bacterial FA composition, we compared the total ion chromatogram of the input 5637 lipid extract against the lipid profile of *L. crispatus* Lc195 grown on that lipid extract. The resulting bacterial FAMEs profile displayed a striking resemblance to the mammalian input profile with 14:0, 16:1 FA complex, 16:0, 18:1 FA complex and 18:0 being the most abundant FA species in both profiles. These data suggest that *L. crispatus* builds its cellular membranes by directly assimilating the available pool of host FAs rather than selecting specific FA species (**Fig. 6C**). Combined with our previous observations, these results highlight that *L. crispatus* membrane FA composition directly reflects the FAs available in the environment, and can shift dramatically depending on media composition, which may result in distinct membrane properties and functional outcomes.

To definitively confirm that the FAs observed in *L. crispatus* membranes were derived from the host epithelial cell lipid extract, we performed a metabolic tracing experiment. We isotopically labeled 5637 bladder epithelial cells by culturing them in [U-^13^C]glucose for two passages to generate isotopically labeled cellular lipids. We then extracted these ^13^C-labeled lipids and supplemented them into Lc195 and Lg637 cultures as the sole FA source. FAMEs GC-MS analysis of *L. crispatus* Lc195 and *L. gasseri* Lg637 grown isotopically labeled host lipids revealed distinct mass shifts in key membrane fatty acids, including oleic acid (18:1.49*cis*), palmitic acid (16:0), and stearic acid (18:0). The detection of these heavy isotopologues in the bacterial FA biomass confirms that FUM lactobacilli can actively scavenge and incorporate FAs from host-derived lipids to fulfill their metabolic requirement for exogenous FAs.

## Discussion

Here we demonstrate that fatty acid (FA) auxotrophy is a universal, conserved metabolic trait of female urogenital microbiome (FUM) *Lactobacillus* species, and that its genetic basis is the loss or functional inactivation of the Type II fatty acid synthesis (FASII) pathway encoded by the *fab* operon. Using stable isotope tracing with [U-¹³C]glucose, we show that prominent FUM species *L. crispatus* and *L. gasseri* incorporate zero glucose-derived carbon into their cellular lipid pools, while generalist Lactobacillaceae retain robust *de novo* FA biosynthetic activity. Comparative genomic analysis of 123 representative Lactobacillaceae species reveals that *fab* operon loss is strongly associated with host-adapted lifestyles and is widespread within the genus *Lactobacillus*, which encompasses the dominant FUM species. In *L. crispatus* lineages that retain the operon, a pervasive frameshift mutation in *fabH*, the gene encoding the rate-limiting initiation condensing enzyme, renders the pathway non-functional in 93.8% of publicly available Fab+ *L. crispatus* genomes. Finally, we demonstrate that FUM lactobacilli can satisfy their FA requirement by direct scavenging of host bladder epithelial lipids, establishing a metabolic basis for the intimate association between these bacteria and the bladder epithelium.

The widespread loss of FASII across FUM *Lactobacillus* species is consistent with the broader principle of adaptive gene loss in host-associated microorganisms. Reductive genome evolution, driven by relaxed purifying selection and the benefits of metabolic offloading in nutrient-replete niches, is well-documented among obligate intracellular bacteria and is increasingly recognized as a defining feature of host-adapted microbiota members under the Black Queen Hypothesis.^2,5^ Within the *Lactobacillaceae* family, genomic reduction has been described previously in niche-restricted species including *L. bulgaricus* and *L. helveticus*, where biosynthetic pathway loss correlates with adaptation to nutrient-rich dairy environments.^51,52^ Our data extend this paradigm to the female urogenital niche and demonstrate that FA biosynthesis represents a dispensable metabolic pathway in this environment. The female vaginal and urinary environments evidently provide sufficient exogenous FAs or lipids to sustain FUM lactobacilli allowing these species to forgo the energetic burden of *de novo* synthesis. Critically, the strong phylogenetic correlation between *fab* operon loss and host-adapted lifestyle observed across the 123-genome Lactobacillaceae dataset argues that this is not stochastic genomic drift but an adaptive response to niche-specific selective pressures, consistent with the well-established principle that biosynthetic gene loss is selectively favored in nutrient-replete environments where the lost metabolite can be reliably scavenged.^2,53^

Our Fab operon synteny analysis reveals two distinct but convergent genetic mechanisms underlying FA auxotrophy in FUM lactobacilli: complete Fab operon loss and inactivation of *fabH* by genetic mutation. Complete operon loss is the dominant mechanism across most FUM species, including *L. gasseri*, *L. jensenii*, *L. johnsonii* and *L. iners*, and represents an irreversible commitment to exogenous lipid dependence. The second mechanism, observed in *L. crispatus* Clades 2 and 3, is particularly informative: a single nucleotide deletion within a homopolymeric poly-adenosine tract introduces a frameshift and premature stop codon in *fabH*, truncating the protein to 210 of its 322 wild-type amino acids and eliminating the C-terminal catalytic core including the essential active site residues His244 and Asn274. The occurrence of this identical mutation in 100% of Clade 2 and Clade 3 Fab(+) *L. crispatus* strains suggests either a strong founder effect or ongoing purifying selection against reversion, consistent with the idea that functional FASII confers no fitness advantage — and may impose a metabolic cost — in the female urogenital niche. The enrichment of Fab(+), functionally competent strains in Clade 1, which is predominantly composed of gastrointestinal isolates, raises the intriguing possibility that FASII is selectively maintained in strains that colonize non-urogentital environments, pointing to a genotype-niche correspondence that merits further investigation.

A key finding of this study is that the FA requirement of FUM lactobacilli is not satisfied by saturated FAs alone: myristic acid (14:0) and PA (16:0) as sole FA sources failed to support growth of *L. crispatus* and *L. gasseri*, while monounsaturated FAs of chain lengths C14 to C20 restored growth with C16 and C18 representing optimal substrates. This requirement perhaps reflects the fundamental role of unsaturated acyl chains in membrane fluidity homeostasis in Gram-positive bacteria.^41^ The bacterial membrane must maintain a critical ratio of unsaturated to saturated phospholipid acyl chains to preserve appropriate bilayer fluidity, and the inability to synthesize the *cis*-double bond through the anaerobic FASII pathway means that an exogenous unsaturated FA source is an absolute requirement for membrane biogenesis.^41^ Notably, the requirement was met equally well by naturally occurring *cis* and *trans* C18 isomers, including elaidic acid (18:1Δ9*trans*) and vaccenic acid (18:1Δ11*trans*), indicating that double-bond geometry per se is less critical than the mere presence of a monounsaturated chain of sufficient length. This promiscuity likely reflects the absence of the stereospecific machinery that would otherwise discriminate between isomers and has practical implications for understanding why Tween-80, which contains both *cis* and *trans* C18 species, is an effective FA supplement. Although saturated FAs cannot serve as the sole FA source, our titration experiments indicate that *L. crispatus* and *L. gasseri* can incorporate and utilize PA when a minimal unsaturated FA fraction is also present, suggesting that the requirement is specifically for the unsaturated component, rather than for total FA carbon.

Our findings complement and significantly extend the recent report by Zhu et al., which demonstrated that *cis*-9-unsaturated long-chain FAs differentially modulate *Lactobacillus* species through species-specific response machinery, namely an oleate hydratase (*ohyA*) and fatty acid efflux pump (*farE*), conserved in non-*L. iners* FUM species but absent in *L. iners*.^37^ Where that study characterized FA response machinery and its therapeutic implications, our work identifies the upstream metabolic context that makes such machinery necessary. FUM lactobacilli require exogenous unsaturated FAs because, unlike their non-FUM counterparts, they lack the capacity for *de novo* FA synthesis, and the *ohyA*/*farE* system can be understood as a mechanism for competing effectively for this obligate nutritional resource in the urogenital environment. Our analysis identified full loss of the syntenic Fab operon in *L. iners* suggesting that it is also a FA auxotroph and similarly depends on Tween-80 as a FA source during growth in MRS. The fact that *L. iners* grows robustly in Tween-80, which is predominately composed of 18:1 species, suggests that the previously observed growth inhibitory effect of 18:1 FAs may be concentration dependent.^37^ The observation that *L. iners* lacks both the FASII pathway and the *ohyA*/*farE* FA response genes while remaining a dominant vaginal colonizer suggests that it also relies on the host environment for FAs but assimilates them through distinct, as yet uncharacterized mechanisms.^35^ Taken together, these two studies establish a central role for FA metabolism in FUM community structure and dynamics that likely shapes the competitive metabolic interactions underlying community state transitions between *L. crispatus*-dominant and *L. iners*-dominant states.

The demonstration that lipid extracts from human bladder epithelial cell lines are sufficient to rescue growth of *L. crispatus* and *L. gasseri* as the sole FA source, and that isotopically labeled host epithelial lipids are directly incorporated into bacterial membranes, provides critical evidence for the physiological relevance of FA auxotrophy in the context of the female urinary tract. The bladder is a nutrient-sparse environment, and the identification of bladder epithelial lipids as a plausible FA source positions FA scavenging as a key adaptive strategy for bladder-resident FUM lactobacilli.^7^ The striking resemblance between the cellular FA profile of *L. crispatus* grown on host lipid extracts and the input mammalian lipid profile indicates that these bacteria do not selectively process or remodel the available FA pool but instead directly assimilate available FA species. This metabolic passivity has an important implication: the membrane FA composition of FUM lactobacilli will vary as a function of the local host lipid environment, potentially generating strain-to-strain and individual-to-individual variation in membrane biophysical properties that could influence adhesion, colonization persistence, and host immune interactions. The finding that host lipid acquisition can satisfy the auxotrophic requirement also raises the question of whether disruption of the host lipid environment, for example, through inflammation, antibiotic-mediated dysbiosis, or hormonal changes could compromise FUM lactobacilli colonization by restricting their FA supply.

The observation that several FUM species, including *L. johnsonii*, *L. jensenii*, *L. vaginalis*, and *L. portuensis*, failed to grow in chemically defined medium (CDM) even with Tween-80 supplementation suggests that FA auxotrophy is only one of multiple nutrient dependencies that characterize these organisms. This is consistent with the precedent set by *L. iners*, which harbors the smallest genome among known *Lactobacillus* species and lacks biosynthetic capacity for multiple amino acids and nucleotide precursors, including a well-characterized cysteine auxotrophy that has been proposed as a therapeutic target.^35^ The pattern emerging across FUM *Lactobacillus* species is one of systematic metabolic streamlining; a convergent reduction in biosynthetic self-sufficiency that mirrors the broader principle of host-adapted bacteria offloading nutrient acquisition to the host environment.^2^ Characterizing the full metabolic dependencies of FUM *Lactobacillus* species through systematic CDM supplementation screens represents an important future direction, both for understanding the ecological constraints that shape FUM community assembly and for the rational formulation of defined culture media that faithfully recapitulate the nutritional environment encountered *in vivo*.

The clinical significance of these findings extends directly to the growing field of live biotherapeutic product (LBP) development for urogenital indications including for UTI prevention. Multiple *L. crispatus*-based LBPs are in active clinical development or have recently completed trials, including LACTIN-V, the VIBRANT multi-strain product, and the Seed Health VS-01 vaginal synbiotic, with colonization efficiency identified as a primary limiting factor in all of these programs.^20,22,54,55^ Our data establish that FA auxotrophy is an intrinsic property of the administered strains that must be satisfied for growth and membrane biogenesis, and that the capacity for engraftment is therefore contingent on the availability of appropriate unsaturated FA species in the target niche. This connects directly to the principle established by Shepherd and Sonnenburg that exogenous strain engraftment in an established microbiota is governed by niche availability, including nutritional niche availability, and implies that the FA composition of the target host environment is a determinant of LBP colonization competence that has not previously been considered in trial design.^32^ In addition, the strict requirement for exogenous unsaturated FAs has practical implications for *in vitro* manufacturing: the FA source used during fermentation and formulation will directly determine the membrane composition of the administered product, potentially influencing its viability during storage, metabolic state upon inoculation, and interaction with the host immune system.

In summary, this study establishes FA auxotrophy as a phylogenetically conserved, genetically encoded metabolic hallmark of FUM *Lactobacillus* species, and identifies the loss or inactivation of the FASII *fab* operon as its molecular basis. Together with prior research, these findings position the healthy FUM microbiome as a community of metabolically reduced, host-dependent organisms whose genomic architecture reflects long-term co-evolution with the female urogenital tract.^35,51,52,56^ The ability to build membranes entirely from host-derived FAs represents a striking example of metabolic synergy between a microorganism and its host, analogous in principle to the nutrient interdependencies documented in obligate endosymbionts, though arising in organisms that retain extracellular, free-living lifestyles.^5^ More broadly, these results argue that understanding the metabolic requirements of FUM lactobacilli and the degree to which the host environment satisfies them is essential for interpreting their ecology, understanding their protective associations with host health, and designing the next generation of microbiome-directed therapeutics for urinary tract infection.

## Materials and Methods

### Bacterial strains and culture conditions

All the bacterial strains used in this study are detailed in **Table S1**. Lactobacilli were cultured under hypoxic conditions of 5% oxygen, 10% carbon dioxide, 85% nitrogen at 37°C using Sheldon manufacturing Bactrox Hypoxia/microaerophilic chamber. De Nisco lab biorepository strains were isolated as part of a previously published study of the postmenopausal urinary microbiome.^8^ Briefly, glycerol stocked urine samples were thawed and plated on multiple solid media types including Blood Aagar, Chromagar Orientation (BD), De Man Rogosa Sharpe, Anaerobic-BAP, Columbia Nalidixic Acid agar and incubated under various atmospheric conditions at 35°C for 4 days as described previously.^8^ Individual colonies were restruck and species identification was performed on well-isolated colonies via PCR and Sanger sequencing of the 16S rRNA gene using the 8F and 1492R universal primers.^57^ Identified isolates were grown in appropriate liquid media for organism (i.e. MRS for lactobacilli) and glycerol stocked. ATCC strains were obtained directly from the American Type Culture Collection (ATCC) and revived in MRS broth (BD Difco^TM^). BEI strains were received directly from BEI Resources Repository and were revived from glycerol stocks on anaerobic blood agar plates (BD BBL^TM^) and incubated in hypoxic conditions for minimum of 48 hours. All experiments were conducted with 24-48hr cultures started from single colonies where individual, well-isolated colonies were inoculated from agar plates into liquid media.

### Mammalian cell culture

5637 (ATCC #HTB-1) and T24 (ATCC #HTB-4) human bladder carcinoma cell lines were obtained from ATCC. These cells were revived in Roswell Park Memorial Institute (RPMI) 1640 medium with glutamine (Sigma-Aldrich) supplemented with 10% fetal bovine serum (FBS, Hyclone) and 2% Penicillin-Streptomycin (Sigma-Aldrich). Cell lines were incubated at 37°C in presence of 5% CO_2_ and 95% air in a humidified incubator. Cell lines were split when they reached 80-90% confluency using a 0.25% trypsin-EDTA solution for detachment.

### End point Growth Assay in mMRS +/- T80

MRS media without Tween-80, termed modified MRS (mMRS), was prepared from constituent parts and was comprised of proteose peptone no.3 yeast extract, ammonium citrate, dipotassium phosphate, magnesium sulfate, manganese sulfate, dextrose, sodium acetate, and beef extract. 24-48hr bacterial cultures were pelleted at 4000 x g for 5 minutes. Cell pellets were washed once in mMRS and resuspended in fresh mMRS broth. The optical density was measured at 600nm using BioTek Synergy H1 microplate Reader and inoculums were normalized to OD_600_ = 0.05 for experimental culture conditions: (1) mMRS without Tween-80 and (2) mMRS containing 0.1% of Tween-80 (equivalent to commercial MRS broth). All cultures were incubated in hypoxic conditions for 48 hours. On completion of incubation, OD_600_ readings were taken on neat culture or on dilutions if OD_600_ readings exceeded 0.8. Readings were blanked using the media controls and multiplied by dilution factor whenever required.

### Fatty Acid Assays in CDM

Chemically defined media (CDM) devoid of any fatty acid supplementation was prepared according to published protocols. ^43^ To test growth on different fatty acids, CDM supplemented 300µM (final concentration) of various fatty acids (FAs) including myristic acid, myristoleic acid, palmitic acid, palmitoleic acid, stearic acid, oleic Acid, elaidic acid, vaccenic acid, eicosenoic acid conjugated on bovine serum albumin (BSA)^58^, or Tween-80 as a control. Experimental conditions involved: (i) CDM without FA, (ii) CDM with 0.01% Polysorbate 80, (iii) CDM with FA and (iv) CDM with BSA vehicle. To set up CDM growth assays, overnight MRS cultures were pelleted and washed once using CDM and resuspended in fresh CDM. Optical density was measured at 600nm and normalized to 0.05 in respective experimental conditions. All the cultures were incubated in hypoxic conditions for 24 hours and OD_600_ measurements were recorded. All readings were blanked using the uninoculated media.

### 13C glucose incorporation assays in CDM

The OD_600_ of overnight MRS cultures were measured and diluted to 0.1 in fresh MRS broth and monitored hourly until they reached mid-log phase (approx. OD_600_ 0.5). The sub-cultures were then pelleted at 9000 x g for 3 mins, washed once with CDM without any glucose. Pellets were resuspended in CDM containing either 1% w/v of ^13^C glucose (Cambridge Isotopes Laboratories Inc.) or 1% w/v ^12^C glucose. OD_600_ was normalized to 0.05 in respective medium and incubated in hypoxic conditions for 24 hours. Post-incubation, cultures were centrifuged at 9000 x g for 5 min, pellets were washed with 0.9% saline. Pellets obtained were flash-frozen in liquid nitrogen, stored at -80°C until lipid extraction and FAMEs analysis.

### Bacterial Lipid Extraction and FAMEs GC-MS

Cultures to be used in FAMEs experiments were pelleted at 9000 x g for 5 mins washed once with 0.9% saline, flash frozen in liquid nitrogen and stored at -80°C for lipid extraction. Frozen bacterial pellets were thawed and resuspended in chilled 80% methanol and transferred to clean glass screw cap tube, Methyl-tert-butyl-ether was added and incubated shaking at room temperature for 1 hour.^59^ Phase separation was induced by adding MS-grade water, vortexing for 15 seconds and centrifugation at 1000 x g for 10 mins. The upper organic phase was collected in new fresh glass tube and a 10:3 v/v MTBE:methanol solution was added to the remaining aqueous phase and incubated for 10 min at room temperature with intermediate vortexing and then centrifuged again at 1000 x g for 10 mins. The resulting organic phase was pooled and stored at -20°C overnight, followed by drying under nitrogen stream. After drying, lipids were resuspended in 1M Methanolic Hydrochloric acid (Sigma) and incubated 80°C for 1 hour. After cooling, hexanes and 0.9% saline were added, samples were vortexed for 1 min and centrifuged at 1500 x g for 10 mins. The top hexanes layer was extracted and transferred to an autosampler vial with an insert for GC/MS and stored at -20°C.

### GC-MS Analysis of Fatty Acid Methyl Esters

Extracted FAMEs were separated and analyzed using an Agilent 7890B Gas Chromatograph coupled to a 5977A single quadrupole mass spectrometer. Separation was achieved on a low-polarity HP-5ms (5% phenyl methyl silox) fused silica capillary column (30m × 0.25 mm i.d., 0.25µm film thickness; Agilent).Samples (1 µL) were injected with a 1:10 split ratio using helium as the carrier gas. The oven program commenced at 100°C with an initial hold for 5 min, followed by a ramp of 3°C/min to a final temperature of 250°C, which was held for 1 min. The post-run oven temperature was set to 40°C, and a 2 min solvent delay was employed to protect the detector. The mass spectrometer was operated in electron ionization (EI) mode using full scan acquisition to characterize and identify FAME species, including saturated (14:0, 16:0, 18:0) and unsaturated (16:1, 18:1 isomers) fatty acids. Data acquisition and subsequent processing were performed using Agilent MassHunter software and custom R script. Identification of FAMEs from samples was done by comparing their retention times on the chromatogram against a Supelco 37 component FAMEs Mix solution (CRM47885, Millipore-sigma).

### Incorporation of ^13^C/12C-glucose into 5637 and T24 bladder epithelial cells

Powdered RPMI 1640 medium with L-glutamine, without glucose and sodium bicarbonate (Sigma) was prepared, pH balanced, sodium bicarbonate was added after adjusting pH to 7 and filter sterilized. Either ^12^C or ^13^C glucose was added to two different aliquots of the reconstituted RPMI 1640 medium supplemented with 5% FBS. The 5637 and T24 bladder epithelial cell lines were passaged in the either ^12^C or ^13^C glucose and allowed to grow until 80-90% confluency. The cells were then passaged for a second time in the same media to achieve higher levels of incorporation of the carbon from glucose on to eukaryotic cell membrane lipids.

### Acidic Bligh-Dyer lipid extraction from 5637 and T24 bladder epithelial cells

Lipids were extracted from 5637 cells using the acidic Bligh-Dyer method.^60^ Bladder epithelial cells were grown to 90-100% confluency, washed two times using Dulbecco’s phosphate-buffered saline (D-PBS) and detached using a sterile cell scraper. Cell suspensions were transferred to pre-weighed 15 mL conical tubes and centrifuged at 1000 x g for 5 min. The supernatant was discarded and cell mass was measured using a analytical scale. Cell pellets were resuspended in PBS and transferred to a pre-weighed glass tube. Chloroform (CHCl_3_) and methanol(MeOH) were added to the resuspended cell pellets in final ratio of CHCl_3_:MeOH:PBS (1:2:0.8 v/v) to form a single-phase Bligh-Dyer. Samples were vortexed for 20 min at room temperature and then centrifuged at 500 x g for 10 mins. Supernatants were collected in a new tube and 37% HCl, CHCl_3_ and PBS was added in the ratio 0.1:1:0.9 respectively to achieve 2-phase Bligh-Dyer. Samples were thoroughly vortexed and centrifuged at 500 x g for 5 mins. The lower phase containing lipids was transferred to a new glass tube and dried under a nitrogen stream.^61^ The mass of the dried lipids was measured before storage at -20°C.

### Supplementing bladder epithelial lipids in mMRS

Approximately 120 mg of dried lipids were resuspended in 200µL of 100% ethanol and 50µL of the resuspended lipids were transferred to a new tube for future GC-MS analysis. The remaining 150µL containing ∼90mg of the lipids were incubated for 2 hours in 9.85 mL of mMRS media free from Tween-80 at a final concentration of ∼9mg/mL, and the media was filter sterilized. Bacterial precultures were centrifuged at 4000 x g for 5 mins and washed once using mMRS media. Pellets were then resuspended in mMRS media and the OD_600_ was measured and normalized to 0.05 in experimental conditions: i) mMRS, ii) mMRS with 5637 lipids, iii) mMRS with T24 lipids. Cultures were incubated for 48 hours in hypoxic conditions. Post incubation, OD_600_ was measured and corrected for absorbance by media controls. Cultures were pelleted at 9000 x g for 5 mins washed once with 0.9% saline, flash frozen in liquid nitrogen and stored at -80°C for lipid extraction.

### Construction and annotation of the family-level Lactobacillaceae phylogenetic tree

Genomes from the_Lactobacillaceae family were collected using the Zheng, 2020 taxonomic note as a guide, as well as complete genomes available on the SyntTax server, prioritizing reference genomes identified by NCBI and genomes with a CheckM completeness of above 90%.^46,62^ At least one genome is present for each of the 31 genera of Lactobacilliaceae (*Lactobacillus, Lacticaseibacillus, Leuconostoc, Companilactobacillus, Fructilactobacillus, Lactiplantibacillus, Apilactobacillus, Lentilactobacillus, Levilactobacillus, Weissella, Limosilactobacillus, Pediococcus, Bombilactobacillus, Ligilactobacillus, Latilactobacillus, Liquorilactobacillus, Paralactobacillus, Convivina, Agrilactobacillus, Dellaglioa, Paucilactobacillus, Fructobacillus, Oenococcus, Furfurilactobacillus, Lapidilactobacillus, Schleiferilactobacillus, Acetilactobacillus, Loigolactobacillus, Amylolactobacillus, Secundilactobacillus,* and *Holzapfelia*). A total of 131 genomes were collected for analysis. ANIclustermap (v1.4.0; https://github.com/moshi4/ANIclustermap) was used to make a dendrogram based on ANI values, computed by FastANI.^63^ The average ANI of the 123 genomes calculated from the clustered all-vs-all ANI matrix output was 86.75. The resulting ANI-based dendrogram was rooted using min-VAR rooting with FastRoot v1.5 and visualized in ITOL.^64,65^ The lifestyle category information was based on Zheng 2020 descriptions of the species, literature search, and the metadata included in the NCBI genome submission.^46^

### Family level Fab Operon Synteny Analysis

Fab operon synteny was tested using the fabF gene as a query on the SyntTax server for all complete genomes available for each analyzed species.^62^ For analysis of contig-level genomes, not available on SyntTax, NCBI BLAST using the *fabF* gene (WP_005715309.1) was performed on each genome assembly to locate the Fab operon, which was then independently validated on Geneious Prime® 2026.0.2. Species were annotated by Fab operon presence with “Present” requiring conservation of all genes essential for type II fatty acid biosynthesis as including *fabH*, *fabk,I* or *V*, *fabD*, f*abG*, *fabF*, *accB*, *fabz*, *accC, accD*, and *accA*. Species were categorized as “Absent” with a white square if their genomes did not have any Fab operon synteny or were missing essential genes. If some genomes of a species had the full Fab operon and other genomes did not have synteny then the species was labeled as “Present in some”, designated with a grey square on the tree.

### Fab operon synteny analysis in urogenital *Limosilactobacillus* species

The *Limosilactobacillus portuensis* and *Limosilactobacillus vaginalis* genome sets were curated from NCBI RefSeq by excluding atypical genomes and metagenome-assembled genomes. Only high-quality genomes with fewer than 200 contigs and greater than 95 CheckM completeness were included in the databases constructed for each species. As of March 17, 2026, 16 genomes were included in the *L. portuensis* database and 35 were included in the *L. vaginalis* database. A tblastn search was performed using WP_005715309.1 as the query on each custom database to identify the *fabF* gene. Manual inspection of the *fabF+* genomes was performed to confirm the presence of the complete Fab operon using Geneious Prime® 2026.0.2.

### Analysis of Fab operon structure conservation across the Lactobacillaceae family

Several species across different Lactobacillaceae genera containing a functionally complete Fab operon were identified using NCBI Prokaryotic Genome Annotation Pipeline annotations in Geneious Prime® 2026.0.2. Thirty-six Fab operon sequences were extracted and exported as a concatenated GFF file. This GFF file was imported into RStudio (R version 4.4.1), converted to a data frame, and the CDS features were extracted. A curated mapping table was used to standardize the original CDS Name fields to a *fab* operon gene name or other category. The order of the operons was determined by an ANI-based dendrogram created with ANIclustermap (v1.4.0; https://github.com/moshi4/ANIclustermap) and aligned using the *fabF* gene of each operon. The comparative plot was generated with the R packages ggplot2 (R package version 4.0.1) and gggenes (R package version 0.6.0), depicting each CDS as a directional arrow. The full R script is available on GitHub (https://github.com/deniscolab/Fab_Operon_Structure_Conservation_Vizualization).

### *Lactobacillus crispatus* species level phylogenetic tree

The *L. crispatus* genome set was curated from NCBI RefSeq by excluding atypical genomes and metagenome-assembled genomes. Only high-quality genomes with fewer than 200 contigs and greater than 74.12 CheckM completeness were included. A total of 299 *L. crispatus* RefSeq genomes available as of April 08, 2025, were included in this study. Two genomes of *L. crispatus* strains isolated from the urine of postmenopausal women sequenced as part of this study (BioProject PRJNA1423329) were also included as well as CTV05, EX849587VC08, and EX533959VC04 genomes that were assembled from publicly available reads (BioProjects PRJNA1423329, PRJNA36325), making the total *L. crispatus* genomes analyzed 304.^14^ ANIclustermap v1.4.0 was used to make a dendrogram based on pairwise average nucleotide identity (ANI) values, calculated by FastANI through an all-vs-all genome comparison. The average ANI of the 304 genomes was 98.2506. The resulting ANI-based dendrogram was rooted using min-VAR rooting with FastRoot v1.5 and visualized in ITOL. Location and Biotype information was derived from NCBI. *L. crispatus* Fab operon presence or absence was determined through gene synteny analysis using the *fabF* gene encoding beta-ketoacyl-ACP synthase II from *Lacticaseibacillus rhamnosus (*WP_005715309.1) as the query protein sequence on the SyntTax web server.^62^ We were able to analyze a total of 27 complete *L. crispatus* genomes on the SyntTax server for *fabF* gene and Fab operon synteny. Because the SyntTax database is limited to complete genomes, we expanded our search by creating a custom nucleotide database of the 299 *L. crispatus* refseq genomes collected (inclusive of the 27 already analyzed via SyntTax) using NCBI BLAST+ v2.16.0. The full nucleotide sequence of the putative Fab operon was extracted from the *L. crispatus* strain FDAARGOS_743, including the following genes: *fabZ, fabT, fabH, acpP, fabD, fabG, fabF, accB, fabZ, accC, accD, accA, fabI*, spanning 9,189 bp. This sequence was used as the query in a BLASTn search of our custom *L. crispatus* database. For genomes analyzed by both Syntax and BLASTn, Fab operon synteny results were identical. In total, 93 genomes (93/299; 31.10%) matched the entire operon query length with identical or near-identical nucleotide identity and were annotated as containing the Fab operon in the ANI-based trees. The same BLASTn method was also used for the 5 genomes (Lc195, Lc1505, CTV05, VC08, and VC04) assembled as part of this work. Lc1505, CTV05, VC08, and VC04 had valid hits that matched the length of the Fab operon, making total number of *L. crispatus* genomes with Fab operon synteny 97/304 (31.91%).

### L. crispatus FabH sequence analysis

Coding sequences of Fab operons identified in *L. crispatus* genomes were manually inspected using Geneious Prime® 2026.0.2, Build 2025-12-05 12:29 to identify any potential mutations or splitting of open reading frames due to premature stop codons. After identification of FabH open reading frame splitting due to a premature stop codon in several *L. crispatus* Fab+ genomes, *fabH* nucleotide and protein coding sequences were extracted from all Fab+ *L. crispatus* genomes, Multiple sequence alignment of the nucleotide sequences was performed using MAFFT (Multiple Alignment using Fast Fourier Transform) v7.526.^66^ The --auto parameter was utilized, allowing the software to automatically select the most appropriate alignment strategy based on the input data size and sequence characteristics. This initial nucleotide alignment was used to identify the precise genomic location of the single-nucleotide deletion. Translation was performed using the standard Bacterial, Archaeal, and Plant Plastid genetic code (Translation Table 11). To fully capture the downstream “scrambled” amino acid sequences and identify premature termination events, the translation algorithm was strictly forced to read continuously without truncating at the first encountered stop codon. The resulting unaligned protein sequences, which contained both the conserved N-terminal regions and the frameshifted C-terminal sequences of the variant strains, were subsequently aligned using MAFFT v7.526 with the --auto parameter.^66^ This approach ensured that homologous wild-type regions aligned correctly, while explicitly demonstrating the complete sequence divergence of the variant strains downstream of the mutation event. In total, 91/97 (93.81%) of the Fab+ *L. crispatus* genomes we analyzed had the *fabH* premature stop. Visual representation of the sequence alignments was generated using pyMSAviz (v0.4.2; https://github.com/moshi4/pyMSAviz), a Python wrapper built upon Matplotlib and Biopython.

### FabH 3D structural prediction and alignment

The amino acid sequences of FabH from *L. crispatus* Indica (wild-type) and LB115 (FabH variant, FabH^var^) strains were submitted to the AlphaFold Server (alphafoldserver.com) for three-dimensional structure prediction using AlphaFold3.^67^ Predicted structures were downloaded in PDB format and imported into PyMOL (Schrödinger, LLC). Structural alignment of the wild-type and FabH^var^ predicted models was performed using the PyMOL align command, which minimizes the root mean square deviation (RMSD) of backbone Cα atoms. The catalytic triad residues His244 and Asn274, previously defined by crystallographic analysis of *E. coli* FabH^49,50^, were mapped onto the wild-type structure and assessed for presence in the FabH^var^ model. Structural figures were rendered in PyMOL.

